# Exoribonuclease RNase R protects Antarctic *Pseudomonas syringae* Lz4W from DNA damage and oxidative stress

**DOI:** 10.1101/2023.06.01.543265

**Authors:** Pragya Mittal, Rashmi Sipani, Apuratha Pandiyan, Shaheen Sulthana, Anurag K Sinha, Ashaq Hussain, Malay K Ray, Theetha L Pavankumar

## Abstract

RNase R is a highly processive, 3’ -5’ exoribonuclease involved in RNA degradation, maturation, and processing in bacteria. In *Pseudomonas syringae* Lz4W, RNase R interacts with RNase E to form the RNA degradosome complex and is essential for growth at low temperature. RNase R is also implicated in general stress response in many bacteria. We show here that the deletion mutant of *rnr* gene (encoding RNase R) of *P. syringae* is highly sensitive to various DNA damaging agents and oxidative stress. RNase R is a multidomain protein comprised of CSD, RNB and S1 domains. We investigated the role of each domain of RNase R and its exoribonuclease activity in nucleic acid damage and oxidative stress response. Our results revealed that the RNB domain alone without its exoribonuclease activity is sufficient to protect against DNA damage and oxidative stress. We also show that the association of RNase R with the degradosome complex is not required for this function. Our study has discovered for the first time a hitherto unknown role of RNase R in protecting *P. syringae* Lz4W against DNA damage and oxidative stress.

**Importance:** Bacterial exoribonucleases play a crucial role in RNA maturation, degradation, quality control and turnover. In this study, we have uncovered a previously unknown role of 3’-5’ exoribonuclease RNase R of *P. syringae* Lz4W in DNA damage and oxidative stress response. Here, we show that neither the exoribonuclease function of RNase R, nor its association with the RNA degradosome complex is essential for this function. Interestingly, in *P. syringae* Lz4W, hydrolytic RNase R exhibits physiological roles similar to phosphorolytic 3’-5’ exoribonuclease PNPase of *E. coli*. Our data suggest that during the course of evolution, mesophilic *E. coli* and psychrotrophic *P. syringae* have apparently swapped these exoribonucleases to adapt to their respective environmental growth conditions.

## Introduction

Exoribonucleases play an important role in determining the levels of RNA in cells (1). Some of the exoribonucleases also associate with the degradosome, a multi-protein complex involved in degradation and processing of cellular RNA (2, 3). The RNR superfamily of enzymes which includes proteins like RNase II, RNase R, and the eukaryotic Rrp44/Dis3 are present in all domains of life (4). RNase R, encoded by the *rnr* gene, is a 3’ to 5’ highly processive exoribonuclease involved in degradation, maturation and processing of RNA in bacteria (5-8). RNase R plays an important role in RNA quality control, such as the degradation of defective tRNA and the removal of aberrant fragments of 16S and 23S rRNAs (7, 9). RNase R is capable of processing dsRNA without the help of a helicase, hence it is implicated in the removal of mRNAs with stable stem loops, including repetitive extragenic palindromic (REPs) elements (6, 10). It is also involved in the processing of tmRNA (transfer-messenger RNA) and is required for the process of trans-translation (11). Structurally, RNase R is a multidomain protein consisting of three important domains: the N-terminal RNA binding cold shock domain (CSD), a central nuclease domain (RNB) and the RNA binding domain (S1) at the C-terminus. (12). The catalytic RNB domain contains four highly conserved Aspartate residues that help to coordinate Mg^2+^ ions in the catalytic center to facilitate ribonucleolytic activity (12-14).

In *Pseudomonas syringae* Lz4W, RNase R associates with endoribonuclease RNase E, along with DEAD box helicase RhlE to form the degradosome complex (15). The *rnr* gene of *P. syringae* Lz4W and *P. putida* is implicated in survival at low temperature (7, 16-18). *P. syringae* Lz4W cells lacking RNase R accumulate unprocessed 3’ ends of 16S rRNAs in polysomes, making the translation machinery inefficient at low temperature, causing cold sensitivity (7). RNase R is also involved in general cellular stress responses. In *E. coli*, RNase R levels increase under cold shock, stationary phase, and heat shock (19, 20).

In a quest to understand the role of *P. syringae* Lz4W RNase R (RNase R^Ps^) in response to general cellular stress, we subjected the *rnr* mutant to various types of stresses and found that *rnr* mutant cells are sensitive to oxidative stress and DNA damaging agents. We show that this role of RNase R is independent of its association with the degradosome complex in *P. syringae* Lz4W. By using a series of domain deletion constructs and mutating catalytically important residues, we discovered that the catalytic RNB domain alone without its exoribonuclease function is sufficient to protect the *rnr* mutant of *P. syringae* from DNA damage and oxidative stress. We hypothesize that either a cryptic function or putative helicase activity of the RNB domain as shown for *E. coli* RNase R, is likely to play a role in protecting *P. syringae* Lz4W cells from DNA damage and oxidative stress.

## Results

### *P. syringae rnr* mutant cells are sensitive to oxidative stress and DNA damaging agents

In an attempt to understand the role of RNase R in general stress response in *P. syringae*, the *rnr* mutant was subjected to various conditions that include low and high temperatures (4 and 37 ºC), osmotic stress (1M NaCl), envelope stress (10% SDS, 100 mg/ml lysozyme with 0.5 mM EDTA), antibiotics treatment (100 µg/ml rifampicin and spectinomycin), oxidative stress (0.25 mM hydrogen peroxide, 15 µM paraquat) and DNA damage (24 J/m^2^ ultraviolet radiation, 0.25 mM cisplatin, 1 µg/ml mitomycin C and 2.5 mM hydroxyurea). As shown in Table 1, we observed that the sensitivity of *rnr* mutant against osmotic, envelope and antibiotic stresses is comparable to that of wild-type (*wt*) cells. As shown earlier, we confirmed that the *rnr* mutant is cold-sensitive (7), nevertheless, we found that the *rnr* mutant is also sensitive to high temperature stress (Table 1).

Interestingly, we discovered that *rnr* mutant cells are highly sensitive to the oxidative stress induced by H_2_O_2_ and paraquat, and to DNA damage induced by ultraviolet (UV) radiation, mitomycin C (MMC) and hydroxyurea (HU) (Table 1). H_2_O_2_ and paraquat generate reactive oxygen species (ROS), such as hydroxyl and superoxide radicals that can easily react with biological macromolecules including DNA, RNA, proteins and lipids (21). We quantified the effect of these agents on the growth and viability of *rnr* mutant in comparison to *wt* cells. We assessed the colony-forming ability of cells upon treatment with various doses of H_2_O_2_, paraquat, UV radiation, MMC and HU. *rnr* mutant cells displayed more than 100-fold higher sensitivity compared to *wt* cells to all the agents tested, suggesting a cryptic role of RNase R in protecting *P. syringae* cells from DNA damage and oxidative stress (Figure 1A). Spot assays on ABM plates containing the indicated amounts of H_2_O_2_, paraquat, MMC, HU and UV irradiation, further confirmed that the *rnr* mutant of *P. syringae* is highly sensitive to all the agents (Figure 1B). In contrast, we observed that mesophilic *E. coli* cells lacking RNase R (Δ*rnr*) are refractory to oxidative stress and DNA damaging agents (Figure S1) suggesting that the protective role of RNase R against DNA damage is specific to *P. syringae*.

**Figure 1.**
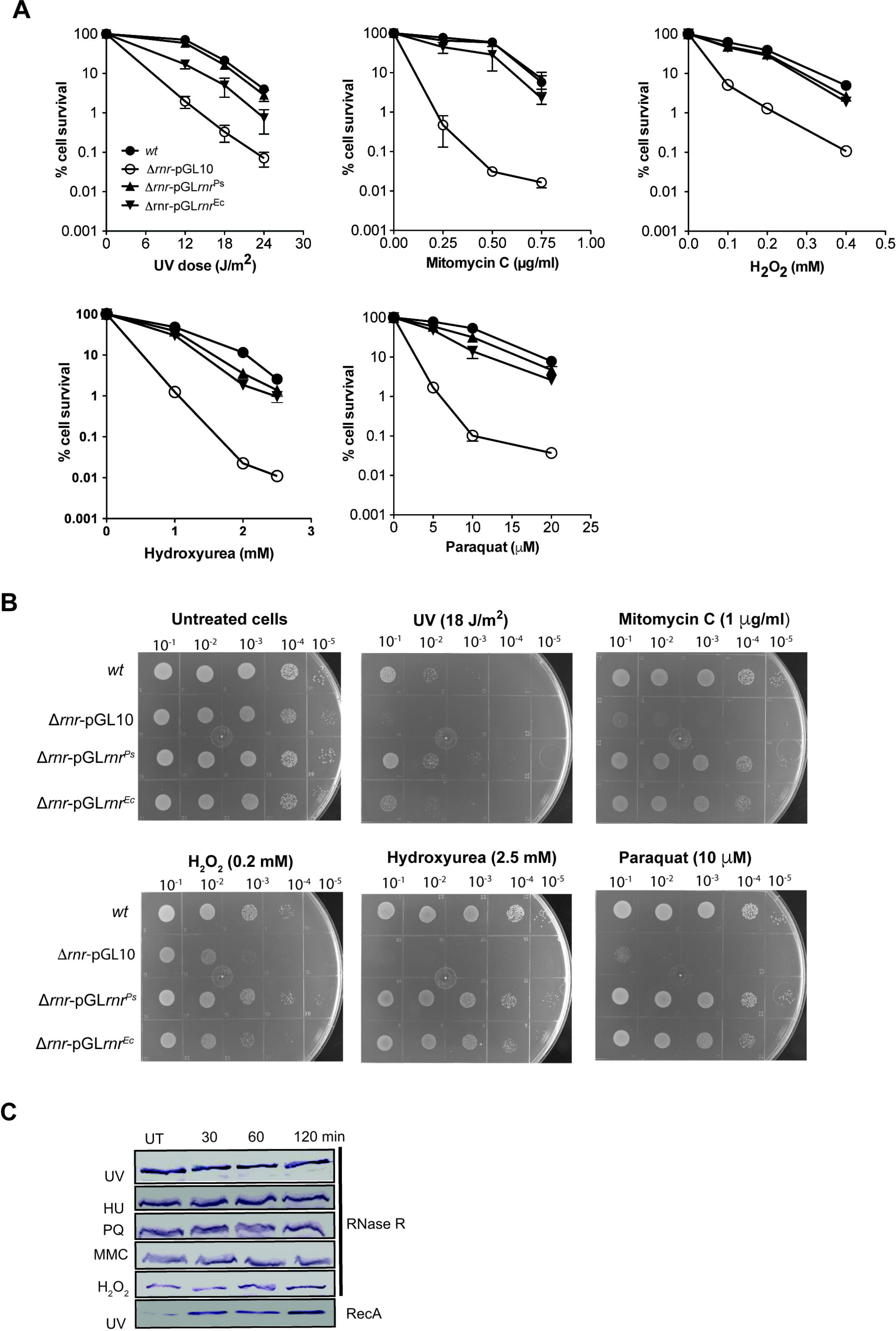
*S*ensitivity of *P. syringae* of Δ*rnr* and complemented strains to DNA damage and oxidative stress causing agents. **A**, Percentage cell survival of WT, Δ*rnr*-pGL10, Δ*rnr*-pGL*rnr*^Ps^ and Δ*rnr*-pGL*rnr*^Ec^ upon treatment with DNA damage and oxidative stress causing agents at indicated concentrations. Data are from the three independent experiments with mean ± SE. **B**, Qualitative assessment of cell viability by spot assays. WT, Δ*rnr*-pGL10, Δ*rnr*-pGL*rnr*^Ps^ and Δ*rnr*-pGL*rnr*^Ec^ were treated with DNA damage and oxidative stress causing agents at indicated concentrations. **C**, Expression levels of RNase R of *P. syringae* upon treatment with DNA damage and oxidative stress causing agents, determined by Western analysis using polyclonal RNase R specific antibodies. The bottom panel shows the expression level of RecA protein upon UV irradiation as a positive control.

Introducing the *rnr* gene encoding RNase R^Ps^ on a low copy plasmid (pGL*rnr*^*Ps-His*^) rescued the growth defects of the *rnr* mutant caused by DNA damage/oxidative stress confirming that the functional loss of RNase R causes the sensitivity (Figure 1A & 1B). Interestingly, when RNase R of *E. coli* (pGL*rnr*^*Ec-His*^) is overexpressed in *P. syringae* Lz4W, it also rescued the growth defect of the *rnr* mutant of *P. syringae* (Figure 1A & 1B). In *E. coli*, another exoribonuclease PNPase has been shown to protect cells against oxidative stress and DNA damage (22-25). We examined if the PNPase of *P. syringae* has similar functions and could complement the loss of RNase R against oxidative and DNA damage stress. However, we observed that the *rnr* mutant over-expressing PNPase could not rescue the sensitivity of *rnr* mutant to oxidative and DNA damage stress (Figure S2).

RNase R is known to be upregulated under different stress conditions in *E. coli* (19, 20), hence we examined the levels of RNase R in *P. syringae* Lz4W cells by Western blotting with polyclonal anti-RNase R antibodies (15). The levels of RNase R remained unchanged after treatment with stress-causing agents (Figure 1C). As a control, the expression of DNA repair protein RecA was induced quickly upon treatment with UV irradiation as shown earlier (26). This finding is consistent with a previous report showing that the RNase R of *P. syringae* Lz4W is not induced under cold stress and in the stationary phase (7). This suggests that, unlike *E. coli*, the RNase R^Ps^ is not a stress-inducible protein.

### RNase R protects *P. syringae* cells against oxidative stress and DNA damage independent of RNA degradosome complex

RNase R associates with endonuclease RNase E and RNA helicase RhlE to form the RNA degradosome complex in *P. syringae* Lz4W (15). The C-terminal domain of RNase E acts as a scaffold for the formation of a degradosome complex with RNase R and RhlE in *P. syringae* (15). Thus, we constructed a C-terminal deletion mutant of RNase E (*rne*^Δ595-1074^) to investigate the role of RNase R associated with the degradosome complex in oxidative stress and DNA damage. Generation of C-terminal deleted *rne* mutant (*rne*^Δ595-1074^) was carried out as described in materials and methods.

We examined RNA degradosome complex formation in the *rne*^Δ595-1074^ strain. We performed the protein fractionation by glycerol density gradient centrifugation as described in the materials and methods. As shown in supplementary figure S3, the co-sedimentation of degradosome proteins RNase E, RNase R wand RhlE proteins was observed only in wildtype cells (Figure S3-C). In contrast, the presence of RNase E, RNase R or RhlE proteins was not detected in the heavier glycerol density gradient fractions (data not shown), suggesting that degradosome complex formation is severely compromised in the *rne*^Δ595-1074^ cells. However, the truncated C-terminally deleted RNase E (RNase E^Δ595-1074^) protein was detected in the precipitated fraction of the cell lysate, suggesting that the solubility of RNase E is greatly affected due to the deletion of the C-terminal region (Figure S3-D), thus affecting the degradosome complex formation.

We further subjected the *rne*^Δ595-1074^ strain to oxidative stress and DNA damage causing agents. Unlike the *rnr* mutant, the *rne*^Δ595-1074^ strain is not sensitive to DNA damage and oxidative stress causing agents and showed resistance similar to *wt* (Figure 2). These results suggest that the requirement of RNase R in protecting *P. syringae* cells against oxidative stress and DNA damage is an intrinsic property of RNase R and is independent of its association with the RNA degradosome complex.

**Figure 2.**
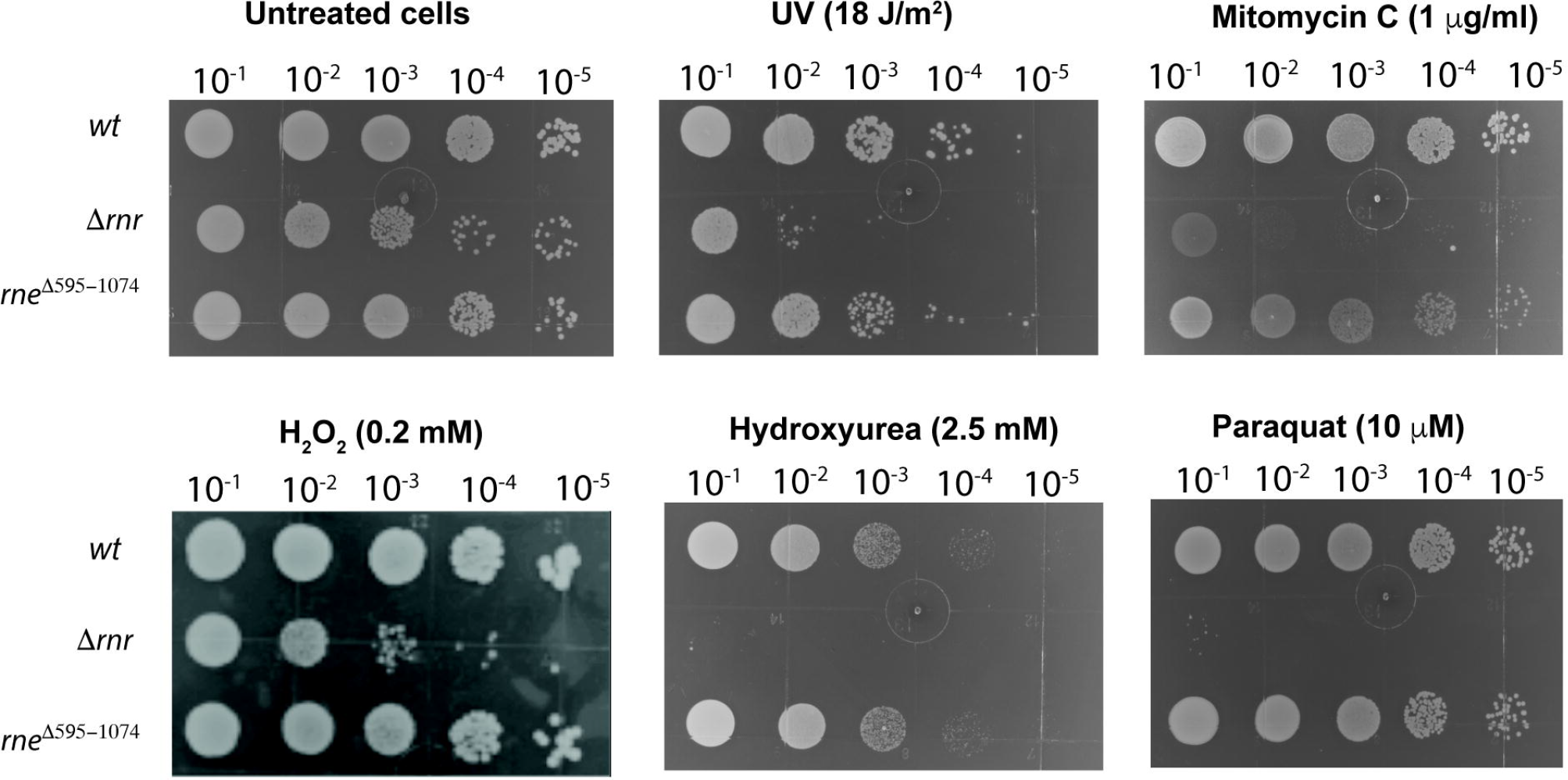
*S*ensitivity of C-terminal domain deletion mutant of *rne* gene (*rne* ^*Δ*595 - 1074^) encoding RNase E compared to Δ*rnr P. syringae* cells. Qualitative assessment of cell viability by spot assays. *wt*, Δ*rnr* and Δ*rne*^Δ595 – 1074^ were exposed to DNA damage and oxidative stress causing agents at indicated concentrations.

### The RNB domain of RNase R is sufficient to protect *P. syringae* Lz4W against oxidative stress and DNA damage

RNase R^Ps^ is a multidomain protein comprising the N-terminal RNA binding cold shock domain (CSD, 1 – 225 aa), the central nuclease domain (RNB, 226 – 665 aa) and the RNA binding domain S1 at the C-terminus (666 – 885 aa) (Figure 3A). To understand the role of each domain in protecting *P. syringae* Lz4W cells against oxidative stress and DNA damage, we constructed a series of domain deletion mutants of RNase R: i) a mutant lacking the CSD domain (ΔCSD), ii) a mutant lacking the S1 domain (ΔS1), and iii) a mutant lacking both the CSD and the S1 domains (RNB) (Figure 3A). In our viability assays, *rnr* mutant cells complemented with domain-deleted mutants of RNase R showed resistance to the oxidative stress and DNA damaging agents similar to *wt* cells (Figure 3B & C). This suggests that the RNB domain of RNase R alone, without the RNA binding CSD and S1 domains is sufficient to protect *P. syringae* Lz4W against oxidative stress and DNA damage.

**Figure 3.**
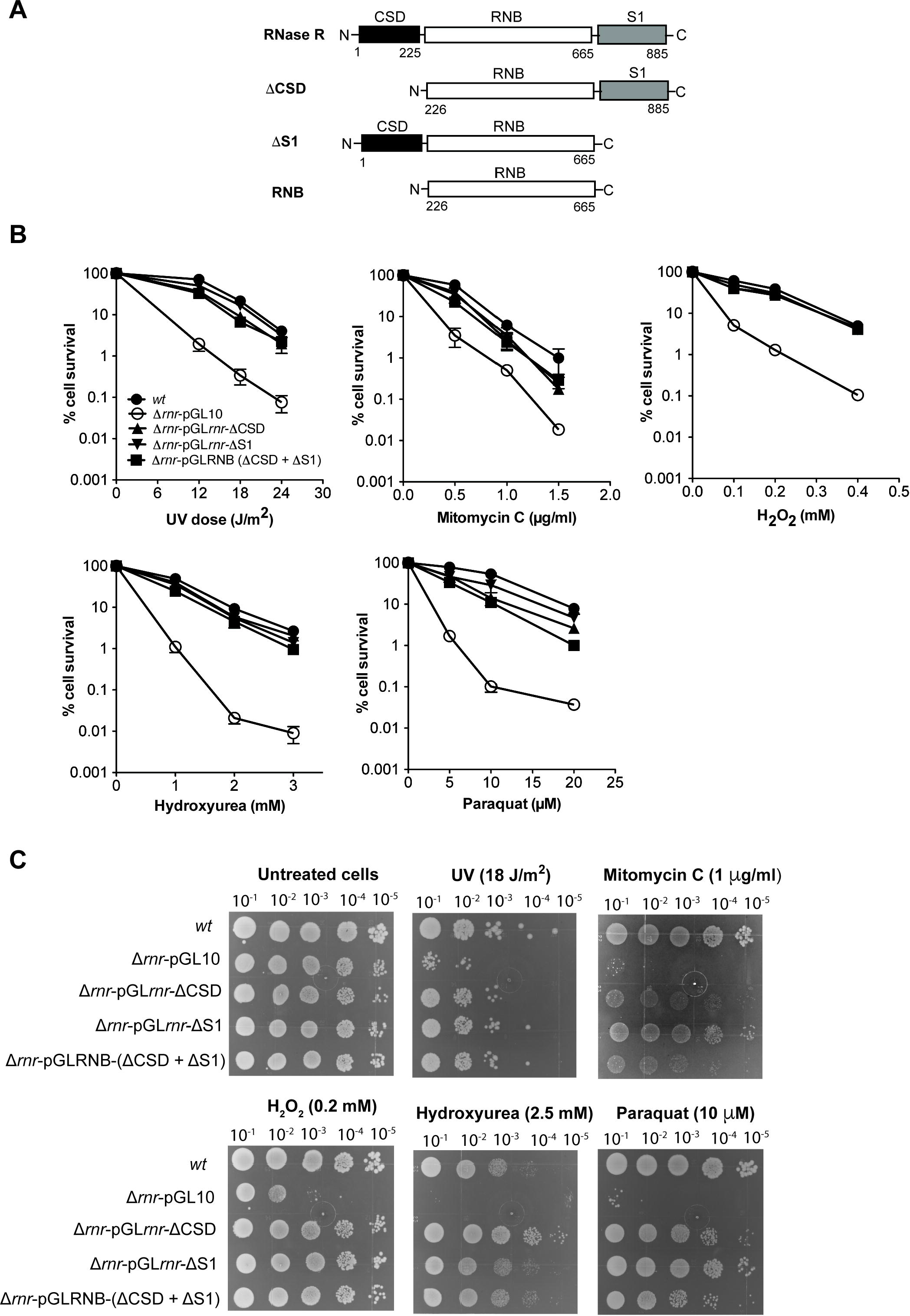
*S*ensitivity of Δ*rnr P. syringae* strain complemented with domain/s deleted mutants of RNase R. A, Schematic representation of structural domains of *P. syringae* RNase R. The domain/s deleted RNase R constructs; cold shock domain (CSD, 1 – 225 aa), catalytic RNB domain (226 – 665 aa), and S1 domain (666 – 885 aa) are shown. **B**, Percentage of cell survival of WT, Δ*rnr*-pGL10, Δ*rnr*-pGL*rnr-ΔCSD*, Δ*rnr*-pGL*rnr-ΔS1* and Δ*rnr*-pGLRNB*(ΔCSD +ΔS1*) upon treatment with DNA damage and oxidative stress causing agents at indicated concentrations. Data are from the three independent experiments with mean ± SE. **C**, Qualitative assessment of cell viability by spot assays. WT, Δ*rnr*-pGL10, Δ*rnr*-pGL*rnr-ΔCSD*, Δ*rnr*-pGL*rnr-ΔS1* and Δ*rnr*-pGLRNB*(ΔCSD +ΔS1*) were treated with DNA damage and oxidative stress causing agents at indicated concentrations.

### Exoribonuclease activity of RNase R is not required for protecting *P. syringae* Lz4W against oxidative stress and DNA damage

The catalytic site of RNase R^Ps^ has four conserved aspartate residues in the active site located at positions 276, 282, 284 and 285 (Figure 4A). D276 and D285 have been proposed to be important for coordinating with Mg^2+^ ion, D282 for holding the RNA in place while D284 for cleaving the phosphodiester bond (27). It was shown in *E. coli* that D280 (equivalent to D284 of *P. syringae*) is the only crucial residue for the exoribonuclease activity of RNase R without affecting its RNA binding (14). This residue is also shown to play a crucial role in the activity of RNase II of *E. coli* and RNase R of *Legionella pneumophila* (28). Hence, we mutated the equivalent Asp284 in RNase R to alanine, overexpressed and purified the mutant protein (RNaseR^D284A^) as described in the materials and methods. The SDS-PAGE profile of RNaseR^D284A^ is shown in Figure 4B(i). RNase R^D284A^ shows double protein bands on a SDS-polyacrylamide gel. The lower band is a C-terminally degraded product of RNase R protein as observed earlier (10). We were unable separate these two bands due to the marginal difference in their molecular weights. We analyzed the exoribonucleolytic activity of RNaseR^D284A^ protein on Poly(A) and *malE-malF* substrates and compared with the wild-type protein. Wild-type RNase R degraded both poly(A) and *malE-malF* substrates within 10 mins (Figure 4B (ii & iii)). Despite using a higher protein concentration of the RNaseR^D284A^ mutant (0.5 µM vs 15 µM), it failed to degrade both substrates even after prolonged incubation. However, a slight degradation product observed at 60 mins, particularly with the *malE-malF* substrate is possibly due to residual activity of the mutant when present at a 30-fold higher concentration than the wild-type enzyme (Figure 4B (ii & iii)). This shows that the D284A mutation leads to a significant loss of exoribonuclease activity of RNase R^Ps^.

**Figure 4.**
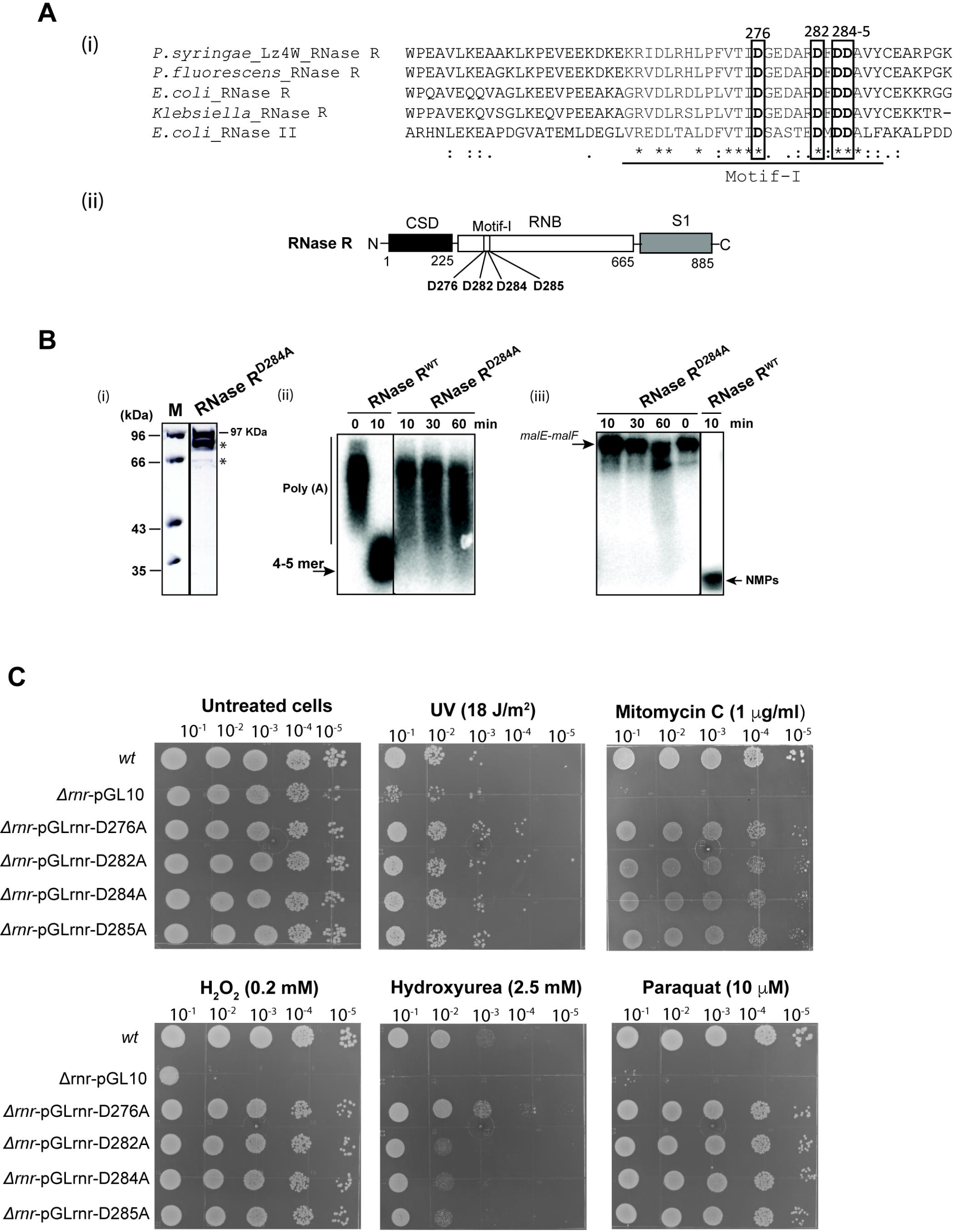
*S*ensitivity of Δ*rnr P. syringae* strain complemented with catalytic site mutants of full-length RNase R to DNA damage and oxidative stress. A, **(i)** The multiple sequence alignment of the catalytic region of *P. syringae* RNase R with RNase R of closely related bacteria including *E. coli*. Boxed and highlighted regions indicate the highly conserved catalytically important aspartate residues in motif-I of catalytic RNB domain of RNase R. **(ii)** Schematic representation of structural domains of *P. syringae* RNase Rand the motif-I of RNB with catalytically important residues are shown. **B**. Purification of RNase R^D284A^ and its exoribonuclease activity on single-stranded and structured RNA substrates. (i), SDS-PAGE analysis of purified RNase R^D284A^, ‘*’ indicate the degraded products of purified RNase R^D284A^ protein. (ii), shows the time-dependent exoribonuclease activity of wild-type RNase R^WT^ and RNase R^D284A^ on single stranded Poly(A) and (iii), *malE-malF* structured RNA substrate at indicated time-points. The end products of size 4-5 mer from degradation of Poly(A), and NMPs (nucleotide monophosphate) from *malE-malF* substrates are indicated. **C**, Qualitative assessment of cell viability by spot assays. The *wt*, Δ*rnr*-pGL10, Δ*rnr*-pGL*rnr-*D276A, *rnr*-pGL*rnr-*D282A, *rnr*-pGL*rnr-*D284A and *rnr*-pGL*rnr-*D285A were exposed to indicated concentrations of DNA damage and oxidative stress causing agents.

In our viability assays, single point mutants of these four highly conserved aspartate residues (D276A, D282A, D284A and D285A) protected Δ*rnr* mutant cells from oxidative and DNA damage stress, similar to *wt* cells (Figure 4C). However, D282A, D284A and D285A mutants showed marginal sensitivity to stress induced by hydroxyurea (HU) suggesting a functional role of these residues in protecting against HU-induced stress. In general, the data suggest that the role of RNase R in protecting cells against oxidative stress and DNA damage is independent of exoribonuclease activity of RNase R.

To further substantiate our finding, we also combined these point mutations in the following combinations: D284A + D285A; D276A + D284A; D276A + D284A + D285A. We examined their functional role in protecting against stress (Figure S4). All double and triple point mutants also protected Δ*rnr* cells against oxidative stress and DNA damage similar to *wt* cells. This further confirms that the exoribonuclease activity of RNase R is not essential for protection against oxidative stress and DNA damage.

### The RNB domain alone with the D284A mutation can protect *P. syringae* Lz4W cells against oxidative stress and DNA damage

The RNB domain of RNase R is the main catalytic domain responsible for its exoribonuclease activity of RNase R. As mentioned above the four conserved Aspartate residues important for the exoribonuclease activity of RNase R are located in motif-1 of the RNB domain (Figure 4A(ii)). To further corroborate our finding that the exoribonuclease function of RNase R^Ps^ is not required for protection against DNA damage and oxidative stress; we mutated the D284 and D284 + D285 residues on catalytic RNB domain alone to alanine. The RNB domain with the D284A and D284A + D285A mutations was expressed in Δ*rnr* strain. Viability assays show that these mutants moderately protect cells against oxidative stress and DNA damage. However, the point mutations slightly affected the ability of the RNB domain to protect cells against mitomycin C, paraquat and hydroxyurea (Figure 5). Taken together, our findings indicate that the RNA degradation function of the RNB domain is not essential for protecting *P. syringae* cells against DNA damage and oxidative stress.

**Figure 5.**
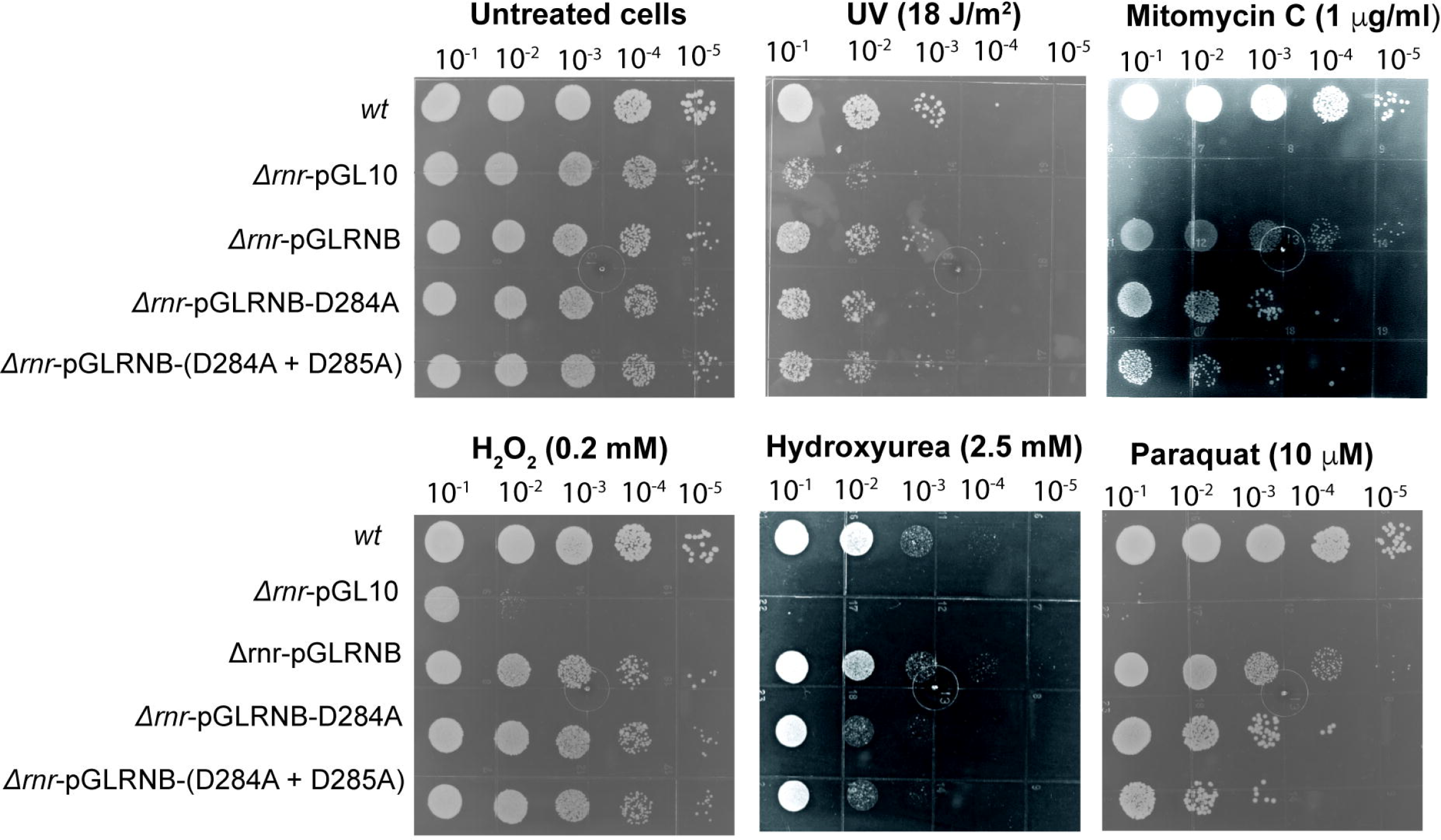
*S*ensitivity of Δ*rnr P. syringae* strain complemented with catalytic site mutants of RNB domain of RNase R. A, Qualitative assessment of cell viability by spot assays. *wt*, Δ*rnr*-pGL10, Δ*rnr*-pGLRNB, Δ*rnr*-pGLRNB-D284A and Δ*rnr*-pGLRNB-D284+D285A were treated with DNA damage and oxidative stress causing agents at indicated concentrations.

In *E. coli*, the RNB domain of RNase R is also suggested to play an important role in RNA unwinding. It has recently been suggested that a triple helix wedge region in the RNB domain of *E. coli* RNase R is responsible for unwinding the duplex-RNA substrates (29). To determine whether RNase R^*Ps*^ possess a similar domain, we compared its predicted structure with the crystal structure of *E. coli* RNase R. Our homology modeling shows that a tri-helix wedge region similar to the one found in *E. coli* RNase R is present in the RNB domain of RNase R^Ps^ (Figure S5), suggesting that the helicase function may be conserved.

## Discussion

Ribonucleases are important modulators involved in various cellular processes by virtue of their RNA processing and degradation activity in maintaining the dynamic transcript levels in bacterial cells. Exoribonucleases like RNase R, PNPase and RNaseJ1 are multifunctional enzymes and are implicated in various biological processes such as stress tolerance, protection from DNA damage and oxidative stress apart from their role in RNA metabolism (19, 20, 22, 24, 30, 31). RNase R is important for the RNA quality control and implicated in various stress conditions in bacteria (5, 20). In this study, we have uncovered a novel role of RNase R in the protection of *P. syringae* Lz4W from oxidative stress and DNA damaging agents such as H_2_O_2_, paraquat, MMC, hydroxyurea and UV radiation.

PNPase is a phosphorolytic 3’-5’ exoribonuclease, an important component of the *E. coli* RNA degradosome complex. However, instead of PNPase, the hydrolytic 3’-5’ exoribonuclease RNase R interacts with the RNA degradosome complex in *P. syringae* (15). Interestingly, the *pnp* mutant of *E. coli* and the *rnr* mutant of *P. syringae* exhibit similar phenotypic behaviors. The *pnp* mutant of *E. coli* and the *rnr* mutant of *P. syringae* are both sensitive to cold, and both enzymes are shown to be involved in the maturation of 3’ end of 16S rRNA (32-35). RNase R of *Pseudomonas putida* is also implicated in survival at low temperature (7, 16-18). PNPase is further implicated in protecting *E. coli* cells against oxidative damage (31) and in the DNA repair process by virtue of its 3’→5’ DNase activity in *B. subtilis* and *E. coli* (22-25).

We show here that the RNase R is involved in oxidative damage and the DNA repair process in *P. syringae* Lz4W. In contrast to *P. syringae* RNase R, the *rnr* mutant of *E. coli* has no discernable phenotype and is refractory to these stress conditions. This implies that the protective role of RNase R is either specific to *P. syringae* Lz4W or possibly specific to *Pseudomonas* species. It is intriguing that although RNase R of *E. coli* is not involved in DNA damage and oxidative stress response in *E. coli*, it protects *P. syringae* Lz4W from DNA damage and oxidative stress. The PNPase^Ec^ and RNase R^Ps^ may have evolutionarily swapped their functions concerning their roles in the cellular milieu of these two bacteria. Although, we don’t know why RNase R has been evolutionary selected over PNPase (unlike *E. coli)* to protect *P. syringae* Lz4W from oxidative stress and DNA damage, we speculate that the highly processive, hydrolytic RNase R is selected over the less processive, phosphorolytic PNPase to conserve energy at low temperature.

RNA degradosome is a multi-enzyme complex that significantly contributes to the steady-state profiles of RNA transcripts in bacteria (36). Our laboratory has previously reported a novel degradosome complex consisting of the RNase E, the exoribonuclease RNase R and a DEAD-box RNA helicase RhlE in *P. syringae* Lz4W (15). We show by deleting the C-terminal domain of RNase E that the association of RNase R with the degradosome complex is not required for protection against oxidative and DNA damage stress, thus suggesting that DNA damage resistance phenotype is an intrinsic function of RNase R.

We further investigated the possible role of each domain of multi-domain RNase R in protecting *P. syringae* cells from oxidative and DNA damage stress. Interestingly, we found the catalytic RNB domain of RNase R^Ps^ alone is sufficient for protecting *P. syringae* cells from oxidative and DNA damage stress, while the CSD domain and the S1 domain are dispensable. In *E. coli*, the RNB domain alone degrades structured RNA (12). Therefore, we envisaged the degradation of structured RNAs either by full-length RNase R^Ps^ or by the catalytic RNB domain itself could play a protective role against oxidative and DNA damage stress in *P. syringae* Lz4W. We replaced the highly conserved aspartate residues of RNase R^Ps^ implicated in its exoribonuclease function with alanine. These mutants showed no apparent sensitivity to oxidative stress and DNA damaging agents, indicating the RNA degradation function of RNase R is not required. Further, the double- and triple-point mutations of the aspartate residues in combinations also resulted in similar results, confirming that the ribonuclease function of RNase R^Ps^ is not essential. Thus, the ability of RNase R to degrade highly stabilized RNA secondary structures does not have a role in protecting *P. syrin*gae cells against DNA damage and oxidative stress. Intriguingly, our data shows that the RNase R^Ps^ lacking exoribonuclease activity and the RNB domain itself can protect cells from oxidative and DNA damage stress, implying the importance of the RNB domain alone with no exoribonuclease activity in this scenario.

If the association of RNase R with the degradosome complex and its exoribonuclease activity is not essential, then how does RNase R protect *P. syringae* cells from DNA damage and oxidative stress? How does a catalytically inactive RNB domain itself without CSD and S1 domains rescue *P. syringae* cells from oxidative and DNA damage stress? Our data suggest that the RNB domain without exoribonuclease function has a cryptic functional role which helps in protecting Lz4W cells from these stress conditions.

RNase R is known to process structured dsRNA without the help of a helicase and is implicated in the removal of mRNAs with stable stem loops (6). It was earlier shown that the cold sensitivity of the *csdA* mutant of *E. coli* can be complemented by overexpressing RNase R, possibly through its helicase function (37). Importantly, the RNB domain of RNase R is also shown to possess helicase activity in *E. coli* (32, 38, 39). A recent study showed the presence of a winged tri-helix domain in *E. coli* RNase R that is responsible for its helicase function (29). This prompted us to investigate the presence of a similar tri-helix wedged domain in RNase R of *P. syringae*. Our homology modeling of *P. syringae* RNase R based on the RNase R of *E. coli*, shows a similar tri-helix wedge in the RNB domain. Hence, we hypothesize that RNase R could participate in the protection of *P. syringae* Lz4W against oxidative and DNA damage stress via its helicase activity. This is further supported by a recent observation in *P. syringae* Lz4W that the DEAD box helicases CsdA and SrmB are involved in protection against DNA damage and oxidative stress (40).

This study uncovered a hitherto unknown role of RNase R in the physiology of cold-adapted bacterium *P. syringae* Lz4W. Here, we report a novel role of RNase R in protecting *P. syringae* Lz4W against DNA damage and oxidative stress. Our study indicates that the protective role of RNase R is specific to *P. syringae* Lz4W. We hypothesize that a cryptic role of RNase R, independent of its exoribonuclease activity, is involved in protecting *P syringae* cells against DNA damage and oxidative stress. In future, it would be of relevance to elucidate the mechanism by which RNase R protects *P. syringae* Lz4W cells from DNA damage and oxidative stress.

## Materials and Methods

### Bacterial strains, plasmids and growth conditions

Bacterial strains and plasmids used in this study are described in Table 2. *Pseudomonas syringae* Lz4W cells were grown in Antarctic Bacterial medium (ABM: 5g liter^-1^ peptone and 2g liter^-1^ yeast extract) at both 22ºC and 4ºC. *E. coli* strains were routinely grown in Luria-Bertani medium at 37ºC. The growth media, when required, were supplemented with antibiotics as follows: 100 µg ml^-1^ ampicillin, 50 µg ml^-1^ kanamycin, 20 µg ml^-1^ tetracycline, and 400 µg ml^-1^ rifampicin. For complementation studies, plasmids were mobilized into *P. syringae* strain by conjugation using *E. coli* S17-1 helper strain. For growth analysis, cultures were grown in ABM overnight and were inoculated in a fresh media in 1:100 dilutions and OD_600_ (O.D at 600 nm) was measured at various time intervals as indicated.

### Spot assays and quantitation of sensitivity to DNA damaging agents

Cells were grown overnight at 22 ºC in AB media containing appropriate antibiotics and inoculated into fresh AB media with 1:100 dilutions. To measure the sensitivity to NaCl (1M), SDS (10% w/v), lysozyme (100 mg/ml) + EDTA (500 mM), ampicillin (100 µg), rifampicin (100 µg), spectinomycin (100 µg), paraquat (20 µM), mitomycin C (1000 ng), H_2_O_2_ (0.2 mM), hydroxyurea (2.5 mM), cisplatin (0.2 mM); up to 5-fold serial dilutions of exponentially grown cells at 22 ºC were made and spotted on ABM agar plates containing stress inducing agents as indicated. Plates were incubated at 22 ºC for 48 h. For UV sensitivity, up to 5-fold serially diluted exponentially grown cultures were spotted on ABM plates and exposed to UV radiations at indicated doses at the rate of 3 J/m^2^/sec and plates were incubated in the dark at 22 ºC for 48 h.

For quantitative analysis, exponentially growing cells were incubated for 30 mins with oxidative stress/DNA damaging agents at indicated concentrations, washed, then serially diluted, and plated on ABM-Agar plates or LB agar plates (for *E. coli* strains). Plates were incubated at 22 ºC for 48 h and the colony forming units (cfu) were scored. The percentage of cell survival was calculated by considering the cfu from untreated cells as 100%.

### Reagents and general DNA recombinant methods

Routine molecular biology techniques like Genomic DNA isolation, restriction enzyme digestion, PCR, ligation, transformation, conjugation, Southern hybridization, Western analysis and RNA isolation were performed as described earlier (41). All restriction enzymes, T4 DNA ligase, T4 Polynucleotide kinase were purchased from New England Biolabs. Oligos used in the study were from Bioserve India (Hyderabad, India). PCR was carried out using proofreading Pfx DNA polymerase from Invitrogen (San Deigo, CA, USA). DNA sequencing was performed using ABI Prism dye terminator cycle sequencing method (Perkin-Elmer) and analyzed on an automated DNA sequencer (ABI model 3700; Applied Biosystems). Antibodies were purchased from Santa Cruz Biotechnology.

For Southern hybridization, genomic DNA was isolated from *wt* and *rne*^Δ595-1074^ mutant cells and digested with PstI enzyme. Products of the restriction digest were separated on a 1% w/v Agarose gel and transferred onto a Hybond N+ membrane (Amersham Biosciences). For the probe, a PCR amplicon of full-length *rne* gene was labeled with [α-^32^P] dATP using a random primer labeling kit (Jonaki, BARC, India). Radioactive signals were detected and quantified by phosphorimager (Fuji FLA 3000).

### Construction of mutants of RNase R and RNB by site directed mutagenesis and genetic complementation studies

The *rnr*^*Ec*^, *rnr*^*Ps*^ genes and the domain deletion mutants of *rnr*^*Ps*^ were amplified from the genomic DNA of Lz4W using the primers listed in Table 3. All constructs contain 6x-Histidine tag at the N-terminal end of the protein. For D276A, D282A, D284A and D285A point-mutations in RNase R^Ps^, site-directed mutagenesis of pGL*rnr*^*Ps-His*^ (7) was carried out using self-complementary primers harboring indicated mutations (listed in Table 3). Subsequently, the double D284A + D285A, D276A + D284 and triple D276A + D284A + D285A mutations were generated by using respective complementary primers using pGL*rnr* ^*Ps-His*^ plasmid with pre-existing mutations. Similarly, the D284A and D284A+D285A mutations were introduced into pGLRNB^-*His*^ vector. Mutations and integrity of full-length genes were confirmed by DNA sequencing.

### Purification of RNase R D284A mutant and biochemical assays

The wild-type RNase R and D284A mutant derivatives were purified by expressing these proteins in the *rnr* null mutant of *P. syringae* Lz4W as described earlier (10). RNase R activity was assayed *in vitro* by its ability to degrade ssRNA substrate Poly(A) and structured RNA substrate *malE-malF* as described earlier (10). Briefly, Poly(A) was labelled at 5’-end using [γ^32^P] and the *malE-malF* substrate was internally labelled using (α^32^P). Assays were carried out in 20-μl reaction mixtures containing 25 mM Tris-HCl (pH 7.0), 100 mM KCl, 0.25 mM MgCl_2_, 5 mM DTT. 10 pmol of labelled substrate was treated with 0.5 µM RNase R^WT^ and 15 µM RNase R^D284A^ at the indicated time-points at room temperature, and stopped by adding 10mM EDTA, 0.2% SDS and 1 mg/ml of proteinase K. RNA degradation products were resolved on 8% denaturing 8 M urea PAGE and degradation products were analyzed by Phosphoimager (Fuji).

### Western analysis

For examining the induction of RNase R, the exponentially grown wild-type *P. syringae* cells were incubated with DNA damaging agents. 1 ml aliquot was collected before treating cells with DNA damaging agents. 5 ml of cells were treated separately with Paraquat (15 µM), MMC (1000 ng), UV (18 J/m^2^), and incubated at 22ºC. 1 ml of aliquot was collected at 30, 60 and 120 min intervals. Cells were further lysed by sonication in the presence of protease inhibitor cocktail (Roche, USA). Protein concentration from cell extracts was measured using Bio-rad Protein Assay kit, also by the absorbance at 280 nm (A_280_) using the Nanodrop spectrophotometer. An equal amount of protein was loaded and resolved on a 10 % w/v SDS-PAGE. The resolved proteins were later electro-blotted onto Hybond C-Extra membrane (Amersham Biosciences) and immunodetection was carried out with polyclonal rabbit Anti-RNase R antibodies as described earlier (15). For detecting the levels of RecA expression, cells were treated with UV (18 J/m^2^) and incubated at 22 ºC in the dark. Culture aliquots were collected after 30 minutes of treatment and immuno-detected by polyclonal anti-RecA rabbit antibodies as described earlier (26).

### Generation of C-terminal deletion of the *rne* gene (encoding RNase E)

For generating C-terminal deletion of *rne* gene, a 784 bp fragment from the N-terminal of *rne* gene was first amplified using primer set Rne_Nterm_FP_BamHI and Rne_Nterm_RP_HindIII having the restriction enzyme sites for BamHI and HindIII respectively (Table 2). A second fragment from the C-terminal of *the rne* gene was PCR amplified using the primer set Rne_Cterm_FP_HindIII and Rne_Nterm_RP_SalI having the sites for restriction enzymes HindIII and SalI (Table 2). Both fragments were initially digested with HindIII and ligated. The ligated product was further digested with BamHI and SalI and cloned into pBlueScript-KS vector. The resultant plasmid pBS(N+C)*rne* was linearized by HindIII, and a tetracycline resistance gene was cloned to obtain the pBS(N+C)*rne*-tet plasmid. The tetracycline resistance gene (*tet*) cassette (∼2.4 KB) was obtained from the pMOS^tet^ (42) as a HindIII fragment. The plasmid construct was confirmed by sequencing and the PCR analysis. Plasmid pBS(N+C)*rne*-tet plasmid was alkaline denatured as described earlier (42). Alkali denatured plasmid DNA was electroporated into the wild-type *P. syringae* cells. Recombinants were selected on ABM plates containing tetracycline and the knock-out was confirmed by Southern and Western analysis.

### Protein fractionation of the RNA degradosome complex

Protein fractionation of the RNA degradosome complex was performed using the glycerol density gradient centrifugation as described earlier (15). Briefly, the cell lysates of wild-type and *rne*^Δ595-1074^ strains were subjected initially to ammonium sulfate precipitation followed by sulfo-propyl Sepharose (SP-Sepharose) chromatography and glycerol density gradient centrifugation. The protein fractions were resolved on an 8 -10 % w/v denaturing polyacrylamide gel and visualized by silver staining the gels. The presence of RNase E, RNase R, and RhlE was identified by the Western analysis using respective polyclonal anti-rabbit antibodies as described earlier (15).

### Modeling of RNase R^*Ps*^ protein

The amino acid sequence of Lz4W RNase R was taken from NCBI (GenBank: AUB77091.1). A model of RNase R^Ps^ was constructed using the SWISS-MODEL (43). The closest template identified was PDB: 5xgu.1, which refers to the RNase R structure from *Escherichia coli*. The RMSD between these two structures was then calculated using the Matchmaker tool of Chimera (44). The RMSD value calculated between 613 pruned atom pairs is 0.248 angstroms; (across all 625 pairs: 0.930). The two structures were superimposed as shown in figure S5.

### Statistical analysis

Data were analyzed using GraphPad Prism version 9.5.0 (GraphPad Software Inc,

USA).

## Acknowledgements

We thank Dr. Manjula Reddy of Centre for Cellular and Molecular Biology, Hyderabad-India for gifting us *wt* and *rnr* mutant cells of *E. coli*. We also thank Drs. Diedre F Reitz and Artem G Lada of Department of Microbiology, University of California, Davis, USA for the comments and suggestions on the manuscript. This study was funded by the Council of Scientific and Industrial Research, New Delhi, India.

