## Supplemental material for "Exoribonuclease RNase R protects Antarctic *Pseudomonas syringae* Lz4W from DNA damage and oxidative stress"

### Supplementary Figures

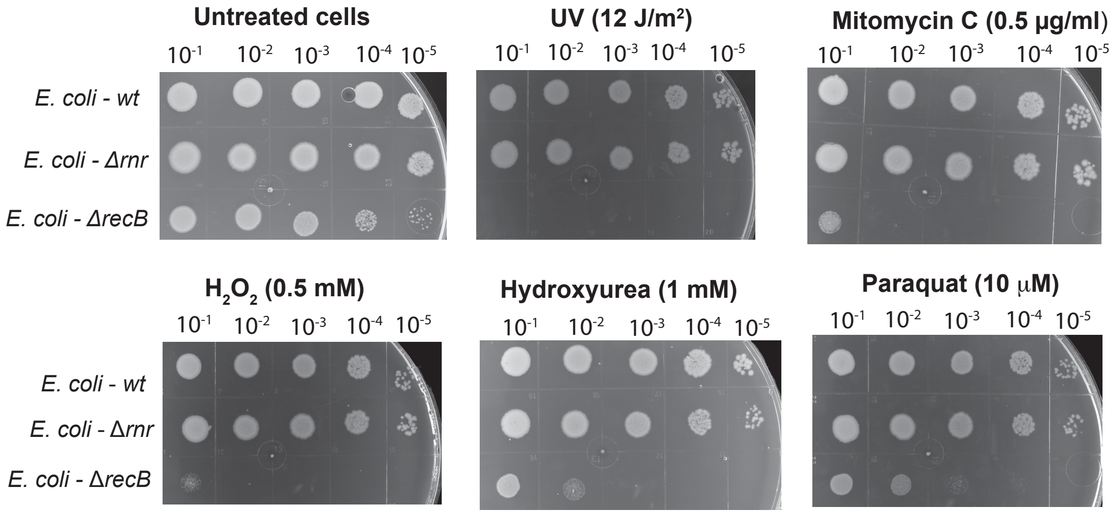


**Figure S1. *S*ensitivity of Δ*rnr E. coli* strain to DNA damage and oxidative stress causing agents.** Qualitative assessment of cell viability by spot assays of *wt*, Δ*rnr* and Δ*recB* mutant of *E. coli*. RecB mutant was used as a positive control against DNA damaging agents. Cells were treated with DNA damage and oxidative stress causing agents at indicated concentrations.

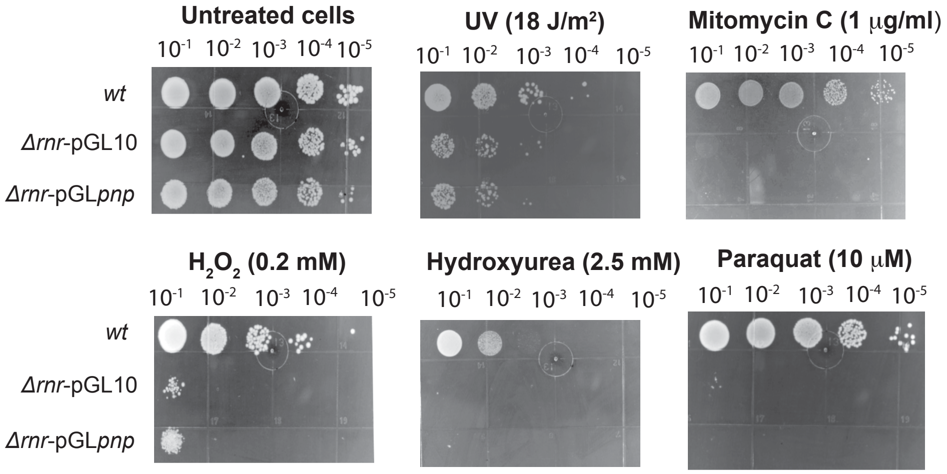


**Figure S2. *S*ensitivity of Δ*rnr P. syringae* strain complemented with PNPase of *P. syringae*.** Qualitative assessment of cell viability by spot assays. *wt*, Δ*rnr*-pGL10 and Δ*rnr*-pGL*pnp* were treated with DNA damage and oxidative stress causing agents at indicated concentrations.

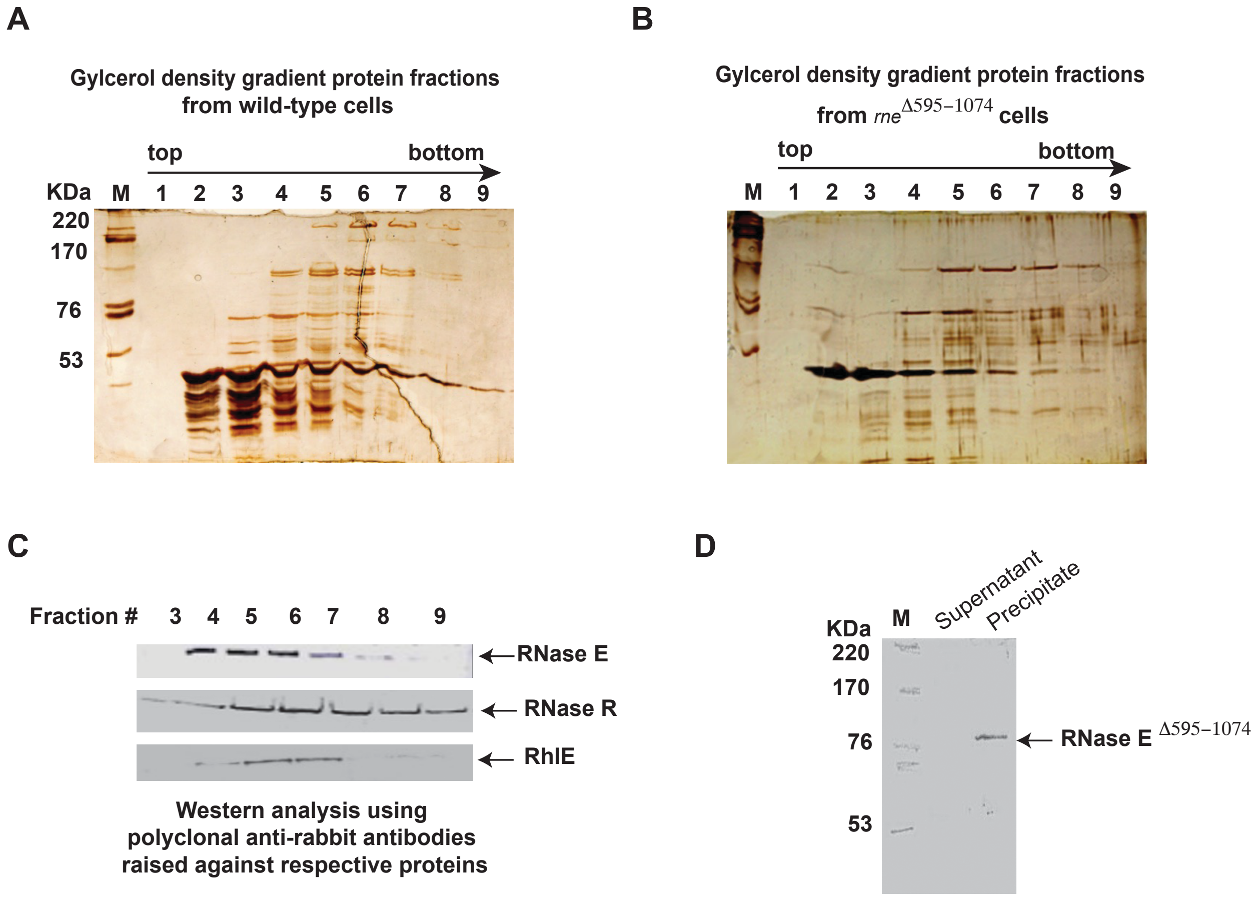


**Figure S3.** ***P. syringae rne***$^{\Delta595\text{-}1074}$ **strain fails to form the RNA degradosome complex.** **A**, the sedimented glycerol density gradient protein fractions (top to bottom; 1 – 9) of wild-type *P. syringae* cells; **B**, the sedimented glycerol density gradient protein fractions (top to bottom; 1 – 9) of the *rne*$^{\Delta595\text{-}1074}$ strain on SDS-PAGE gels with silver staining. **C**, the RNase E, RNase R, and RhlE proteins identified by their specific anti-rabbit polyclonal antibodies from the glycerol density gradient protein fractions of wild-type cells. **D**, the RNase E$^{\Delta595\text{-}1074}$ identified by RNase E specific anti-rabbit polyclonal antibodies from the precipitated fraction of *rne*$^{\Delta595\text{-}1074}$ cells.

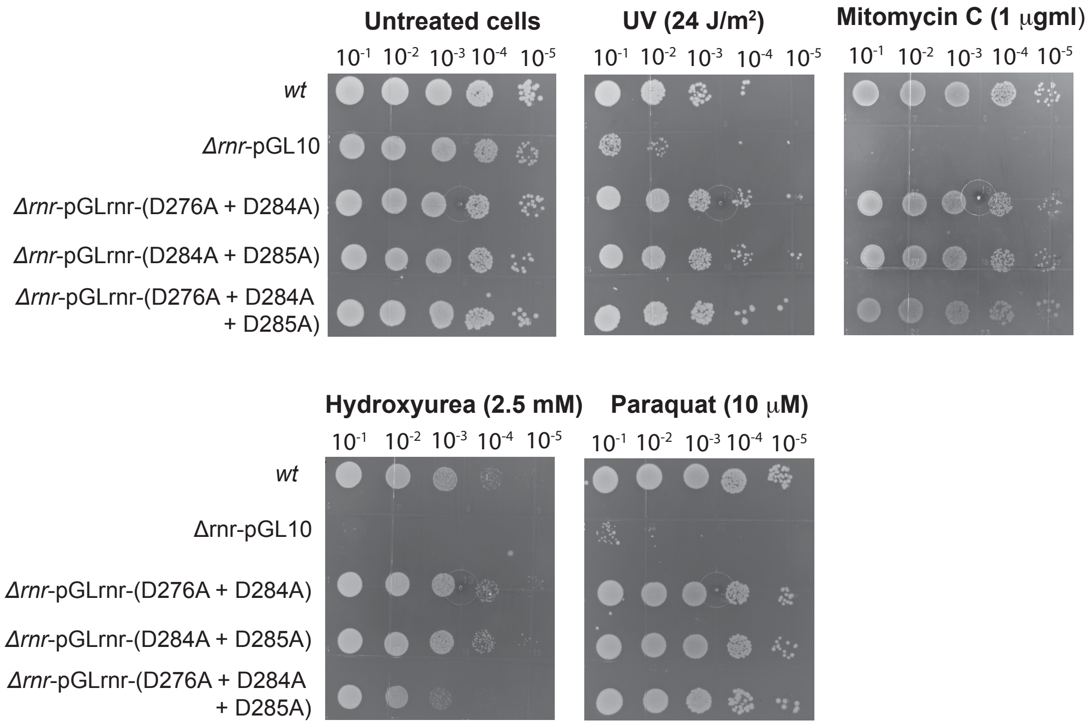


**Figure S4. *S*ensitivity of Δ*rnr P. syringae* strain complemented with double- and triple-mutants of catalytic site residues of RNase R.** Qualitative assessment of cell viability by spot assays. *wt*, Δ*rnr*-pGL10, *rnr*- pGL*rnr*-(D276A+D284A), *rnr*- pGL*rnr*-(D276A+D285A) and *rnr*-pGL*rnr*-(D276A+D284A+D285A) were treated with DNA damage and oxidative stress causing agents at indicated concentrations.

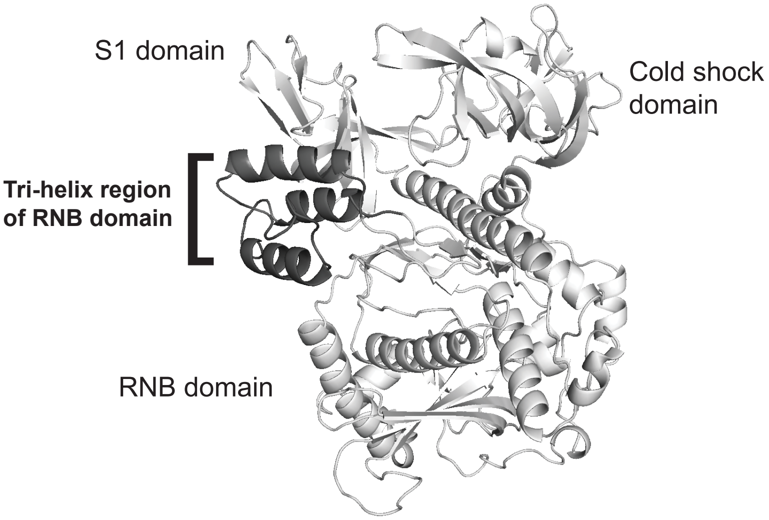


**Figure S5. A structural model of *P. syringae* RNase R constructed by superimposing on the structure of *E. coli* RNase R (PDB:5xgu.1) using SWISS-MODEL.** The tri-helix wedge region of the RNB domain of *P. syringae* RNase R is shown in black color.
